# CD4+ T cells facilitate replication of primary HIV-1 strains in macrophages and formation of macrophage internal virus-containing compartments

**DOI:** 10.1101/2024.03.22.586250

**Authors:** Sabina Victoria Montero, Johanna Leyens, Lea Marie Meckes, Gabriela Turk, Michael Schindler

## Abstract

HIV-1 infects CD4+ T cells and macrophages. However, replication of HIV-1 in these cell types is highly variable and may depend on the use of CCR5 as a co-receptor. In addition, there is internal accumulation of infectious HIV-1 in so-called virus-containing compartments of macrophages (VCCs). VCCs are thought to represent a persistent viral reservoir that is shielded from the antiviral immune response. To date, VCC formation has only been studied in lab-adapted HIV-1 and it is unknown whether VCCs play a role in the replication of primary HIV-1 strains. Furthermore, although macrophages transmit HIV-1 from VCCs to CD4+ T cells, it is unknown whether T cells have an impact on VCC formation. We analyzed the ability of primary and lab-adapted HIV-1 to replicate in macrophages, the effect of coculture with non-infected CD4+ T cells and the extent of VCC formation. Although differentially, all HIV-1 strains replicated in CD4+ T cells, whereas only lab-adapted HIV-1 replicated in macrophages. Strikingly, replication of patient-derived HIV-1 in macrophages was enhanced by coculture with non-infected CD4+ T cells and correlated with VCC formation. In conclusion, non-infected CD4+ T cells facilitate the replication of primary HIV-1 strains in macrophages and the formation of VCCs appears to be a proxy for this phenotype. Our study suggests an essential role for VCCs in the replication of patient-derived HIV-1 in macrophages, which is fueled by non-infected CD4+ T cells. Furthermore, our findings call for strategies to specifically disrupt VCC formation in order to eliminate the HIV-1 reservoir in macrophages.

**IMPORTANCE:** Here we focus on the intimate interplay between HIV-1 infected macrophages and CD4+ T cells. Specifically, we analyzed whether primary HIV-1 strains induce virus-containing compartments (VCCs) within macrophages, which are thought to serve as viral sanctuaries and macrophage reservoirs. Notably, primary HIV-1 strains were unable to replicate in macrophages and induce VCCs unless they were cocultured with non-infected CD4+ T cells, leading to increased VCC formation and viral replication. This suggests an essential role for non-infected CD4+ T cells in facilitating primary HIV-1 replication in macrophages. Our data highlight the importance of not only targeting the latent HIV-1 T-cell reservoir, but also targeting VCC formation in macrophages to achieve the ultimate goal of functional HIV-1 cure.

## INTRODUCTION

Human immunodeficiency virus (HIV) has spread worldwide since its first identification as the causative agent of acquired immune deficiency syndrome (AIDS) in 1983 (1). More than 85 million people have been infected with HIV and 40 million people have died worldwide due to AIDS-related illnesses (2). HIV mainly infects CD4+ T cells, monocytes, macrophages and dendritic cells since these cells express the CD4 receptor, as well as the co-receptors CCR5 or CXCR4 that are required for its entry into target cells (3). Upon prolonged HIV-1 infection there is a drastic depletion of infected and activated CD4+ T cells which leads to the loss of cell-mediated immunity, rendering the patient susceptible to fatal opportunistic infections that is defined as AIDS.

Global scientific efforts to combat AIDS have led to the development of efficient antiretroviral therapy (ART) regimens that suppress viral replication by targeting various steps of the viral life cycle. By decreasing the plasma viral load in patients and preventing disease progression, antiretroviral therapy restores the life expectancy of the patients to an almost normal level (4, 5). Although ART is highly beneficial in controlling HIV infection, it requires lifelong administration and has various side effects. To date, a HIV vaccine could not be developed and there is no cure available that can eradicate the virus from an infected individual. Therefore, there is an ongoing need to understand HIV-1 persistence and transmission in order to devise novel prophylactic and therapeutic treatment strategies (6).

Macrophages are known to be among the first cells that encounter HIV-1 during sexual transmission. They phagocytose pathogens or particles in submucosal tissues and transport them to draining lympnodes, where they present the antigens to CD4+ T cells and activate CD8+ cytotoxic T-cells (CTL), initiating immune control of the infection (7, 8).

Upon the interaction of a CD4+ T cell with an antigen presenting cell (APC), adhesion molecules on the surface of CD4+ T cells probe the APC surface. Recognition of antigen peptide/MHC-II complex leads to the formation of an immunological synapse by concentration of T cell receptors, CD4 and other signaling molecules. This event stabilizes the cell contact which allows T cell activation and proliferation signals to be sustained (9).

Macrophages can be productively infected by HIV-1 and represent an important reservoir even in patients under ART (8, 10, 11). They sequester the virus within an endosomal compartment that is protected from the extracellular milieu (12, 13). These HIV-1-containing compartments in macrophages (VCCs, virus-containing compartments) were shown by EM analysis to be composed of invaginated plasma membrane folds that might remain connected to the cell surface (12, 14, 15). HIV-1 can be rapidly translocated from macrophage internal VCCs via the so-called virological synapse to infect T cells (16).

Most studies that investigate HIV transmission and pathogenesis take advantage of the availability and convenience of laboratory adapted HIV-1 strains. Although they are easy to employ as they efficiently replicate in cell lines, lab-adapted HIV-1 might have lost important features of primary HIV-1 strains. On the other hand, primary HIV-1 strains more closely resemble HIV-1 infection in patients since they are directly isolated from plasma specimens. During more than 80% of the sexual transmission events of HIV-1, a single virus is transmitted from the donor to the recipient. This virus establishes the infection in the recipient and the virus isolated from a patient during the first six months of the infection is therefore called a transmitted/founder (T/F) virus. As the infection progresses to the chronic stages, the virus population in the blood diversifies due to mutations and selection pressure caused by the adaptive immune responses of the host. At the chronic stage, individuals carry HIV-1 populations that are diversified and the virus isolated from the patient’s blood at this period is called a chronic virus (17).

Up to now, studies analyzing the transmission of T/F and chronic HIV-1 strains from myeloid to CD4+ T cells are sparse (18–20) and it is required to further study modes of primary HIV-1 replication and transmission in *in vivo* relevant target cells. Furthermore, investigation of HIV-1 VCC formation and involvement of these structures in transmission from HIV-1 infected macrophages to CD4+ T cells employed laboratory-adapted viral strains such as HIV-1 NL4-3 or AD8. Here, we addressed these shortcomings by employing a set of T/F and chronic HIV-1 strains. We analyzed how these isolates infect primary macrophages and the impact of autologous CD4+ T cell coculture on this. Additionally, we investigated the efficiency of VCC formation by T/F and chronic HIV-1 isolates.

## MATERIAL AND METHODS

### Cell lines

HEK-293T cells (DSMZ ACC635) cells were maintained in Dulbecco’s modified Eagle’s medium (DMEM, high glucose, GlutaMAX™ supplement, Gibco) supplemented with 10% heat-inactivated fetal calf serum (FCS; Invitrogen), 100 µg/ml streptomycin and 100 units/ml penicillin (Sigma) and were cultured at 37°C, 90% humidity and 5% CO_2_.

### HIV-1 constructs

Infectious molecular clones pHIV-1 AD8, pYK-JRCSF, pBR-NL43-V3 92th014.12 have been described (15), as well as pHIV-1 CH077, pHIV-1 THRO, pHIV-1 CHO58 (21) and pUC57rev_HIV-1 C CH293 (17).

### Generation of virus stocks

To produce HIV-1 stocks, 0.5 x 10^6^ HEK-293T cells were seeded per well in a six-well plate and 1 day later the cells were transfected using the calcium phosphate method with 5 µg of the respective HIV-1 molecular clone or mock transfected. At 16 h post transfection, medium was changed and supernatants containing the HIV-1 stock were collected 24 h later and cleared by centrifugation at 3200g at 4°C for 10 minutes.

To generate VPX-VLPs, the above-mentioned protocol was followed and HEK-293T cells were transfected with 5 µg of pSIV_Vpx1 (22) and cotransfected with 0.5 µg of pHIT-G (vesicular stomatitis virus G protein, VSV-G). At 16 h post transfection, medium was changed and supernatants containing the VLP stock were collected and cleared 24 h later.

### Isolation and differentiation of primary human macrophages

Macrophages were generated from buffy coats following established protocols (15, 23). Briefly, peripheral blood mononuclear cells (PBMCs) were isolated from buffy coats by Ficoll-Paque™ PLUS (GE) density-gradient centrifugation. Subsequently, 2.0×10^7^ PBMCs were seeded in petri dishes (Greiner Bio-One) in RPMI 1640 (RPMI-1640 (GlutaMAX™ Supplement; Gibco) medium supplemented with 4% human AB serum (Sigma), 2 mM L-glutamine (PAA), 100 µg/ml streptomycin and 100 units/ml penicillin (Sigma), 1 mM sodium pyruvate (Gibco), 1× non-essential amino acids (Invitrogen) and 0.4× minimal essential medium [MEM] vitamins (Biochrom). Monocytes were differentiated for 3 days by plastic adherence into monocyte-derived macrophages (MDM). Following this initial differentiation phase, non-adherent cells were removed by washing and the MDM were further differentiated for 4 days. Accutase® (Sigma) was used to detach the MDM for 45 minutes at 37°C. The use of PBMCs from healthy human donors was approved by the ethics committee of the University Hospital Tübingen (507/2017BO1, 127/2022BO2 and 860/2023BO2).

### Isolation and maintenance of CD4+ T cells

To isolate CD4+ T cells from PBMCs, the RosetteSep Human CD4+ T Cell Enrichment Cocktail (StemCell) was used according to the manufacturer’s instructions. CD4+ T cells were cultured in RPMI-1640 (GlutaMAX™ Supplement; Gibco) medium supplemented with 10% FCS (Invitrogen), 100 µg/ml streptomycin and 100 U/ml penicillin (Sigma), 10 ng/ml interleukin-2 (IL-2) (Stemcell). Cells were maintained under standard conditions at 37°C, 90% humidity and 5% CO_2_. Following isolation, the maintenance and expansion of T cells was done by adding medium with 10 ng/ml interleukin-2 (IL-2, Stemcell) (10 ng/ml) and for activation Lectin from Phaseolus vulgaris (red kidney bean, PHA) (Sigma, 1µg/ml).

### MDM and CD4+ T cell infection

For MDM and CD4+ T cell infection, HIV-1 stocks were quantified by p24 enzyme-linked immunosorbent assay (ELISA) (24) immediately after virus stock collection. MDM were seeded (5000 per well) in a 96-well plate (Agilent, clear flat bottom), 24 h later, cells were transduced with VPX-containing VLPs for 2 h before infection. Primary CD4+ T cells pre-stimulated with PHA (Sigma, 1µg/ml) and 10 ng/ml IL-2 (Stemcelll) were seeded (200000 cells per well) in a 96-well plate, U bottom (Greiner). Subsequently, the cells were infected for 16 h (for MDM) and 4-6 h (for CD4+ T cells) with 125, 250 and 500 ng/mL of p24 of the HIV-1 strains mentioned in “HIV-1 constructs and generation of HIV-1 stocks”. After 3 dpi the MDM media was replenished and at 7 dpi (for MDM) and 4 dpi (for CD4+ T cells) the cells were washed with PBS and fixed with 2% paraformaldehyde (PFA) for 10 minutes at RT and subjected to intracellular p24 staining. Additionally, samples of the infected cell culture supernatants were collected to quantify virus production by p24 ELISA.

### MDM and CD4+ T cells infection in cocultures

MDM were seeded (5000 per well) in a 96-well plate (Agilent, clear flat bottom). 24 h later, cells were transduced with VPX-containing VLPs for 2 h before infection. Subsequently, the cells were infected for 16 h with 125, 250 and 500 ng/mL of p24 of the respective HIV-1 strains. After 3 dpi, CD4+ T cells pre-stimulated with PHA and IL-2 were added to the macrophages culture at a 1:10 ratio (1 MDM: 10 CD4+ T cells). HIV-1 replication was analyzed in CD4+ T cells and in the MDM as explained below at 8 dpi.

### Quantification of HIV-1 replication in MDM

Upon removal of the CD4+ T cells in the cell culture supernatants the adherent MDM in the cultures were extensively washed with PBS to remove any remaining T cells. After that, the MDM were fixed with 2% PFA for 10 min at 37°C, permeabilization was performed with 1% Saponin-PBS for 10 min at RT, and blocked with PBS + 10% FCS for 20 minutes. Then, cells were stained with primary anti-HIV-Gag antibody (Beckman-Coulter; KC57-RD1, mouse derived), and with α-mouse Alexa Fluor555-conjugated secondary antibody (Invitrogen; goat derived). DAPI was used for nuclear staining and to control for remaining T cells in the microscopical analysis. Finally, cells were washed and fixed with 2% PFA. Fluorescence microscopy images (4X and 40X magnification) were acquired with the Cytation™ 3 Cell Imaging Multi-Mode Reader (BioTek® Instruments). Macrophage-residing HIV-1 in VCCs was analyzed with 40X magnification images. Filters used were the following: “Blue filter“: LED 405 nm, Emissions maximum/bandwidth: 460/60 nm. “Green filter“: LED 505 nm, emissions maximum/ bandwidth: 542/27 nm; “Red filter“: LED: 590 nm, emissions maximum/ bandwidth: 647/57 nm. Images were analyzed using Gen5 Software (version 5.3.10).

### Quantification of HIV-1 replication in CD4+ T cells

After the CD4+ T cells were fixed with 2% PFA for 10 min at 37°C, permeabilization was performed with ice-cold 90% Methanol in H_2_O for 20 minutes on ice, following a blocking step with PBS + 10% FCS for 20 minutes. Then, cells were stained with primary anti-HIV-Gag antibody (Beckman-Coulter; KC57-FITC, mouse derived), and cells were acquired using the MACs Quant Flow Cytometer (Milentyi) equipped with a 488 nm laser and a band filter (525/50). For the data analysis Flow Logic (Flow Logic^TM^) software was used.

### Statistical analyzes

All statistical calculations were performed using GraphPad Prism (version 9.4.1). Unless otherwise stated, data are shown as the mean of at least three independent experiments ± SD. Simple linear regression analysis was performed for the correlation studies and the Pearson correlation coefficients as well as p-values are shown.

## RESULTS

### CD4+ T cells but not macrophages support the replication of primary patient-derived HIV-1 strains

We first aimed to analyze basic characteristics of virus-infection and replication of varying HIV-1 strains in CD4+ T cells and macrophages. For this, lab adapted strain AD8, which is macrophage tropic (25, 26), as well as an CCR5-tropic version of NL4-3 (15, 27) were employed. Furthermore, we utilized the CCR5-using JRCSF, that was isolated from cerebrospinal fluid of a chronic HIV-1 patient and classified as low- or non-macrophage tropic (25, 28) as well as four different primary HIV-1 strains that are all capable of using CCR5 as entry receptor (17, 21). For a robust readout, macrophages were pre-treated for two hours with SIV Vpx-VLPs (29, 30). This inactivates the known HIV-1 restriction imposed by SAMHD1 (31) allowing the assessment of SAMHD1-independent mechanisms that may affect the infection of macrophages by primary patient-derived HIV-1. Then, macrophages were infected with the different HIV-1 strains using increasing amounts of virus normalized to the amount of p24 and the efficiency of virus replication and spread in the culture was assessed 7 days later by p24-immunofluorescence (IF)-staining to identify infected cells (% p24+/DAPI+). While the lab-adapted R5-utilizing AD8 (∼25% p24+ cells) and NL4-3 (∼15% infected cells) showed a high proportion of infected cells at that time point, as expected, the primary HIV-1 strains were severely impaired in their capability to cause efficient infection and spread (less than 1% of infected cells, compare Fig. 1A). Only the primary isolate CH058 showed evidence of low-level infection and replication, with 2-4% of infected macrophages.

**Figure 1.**
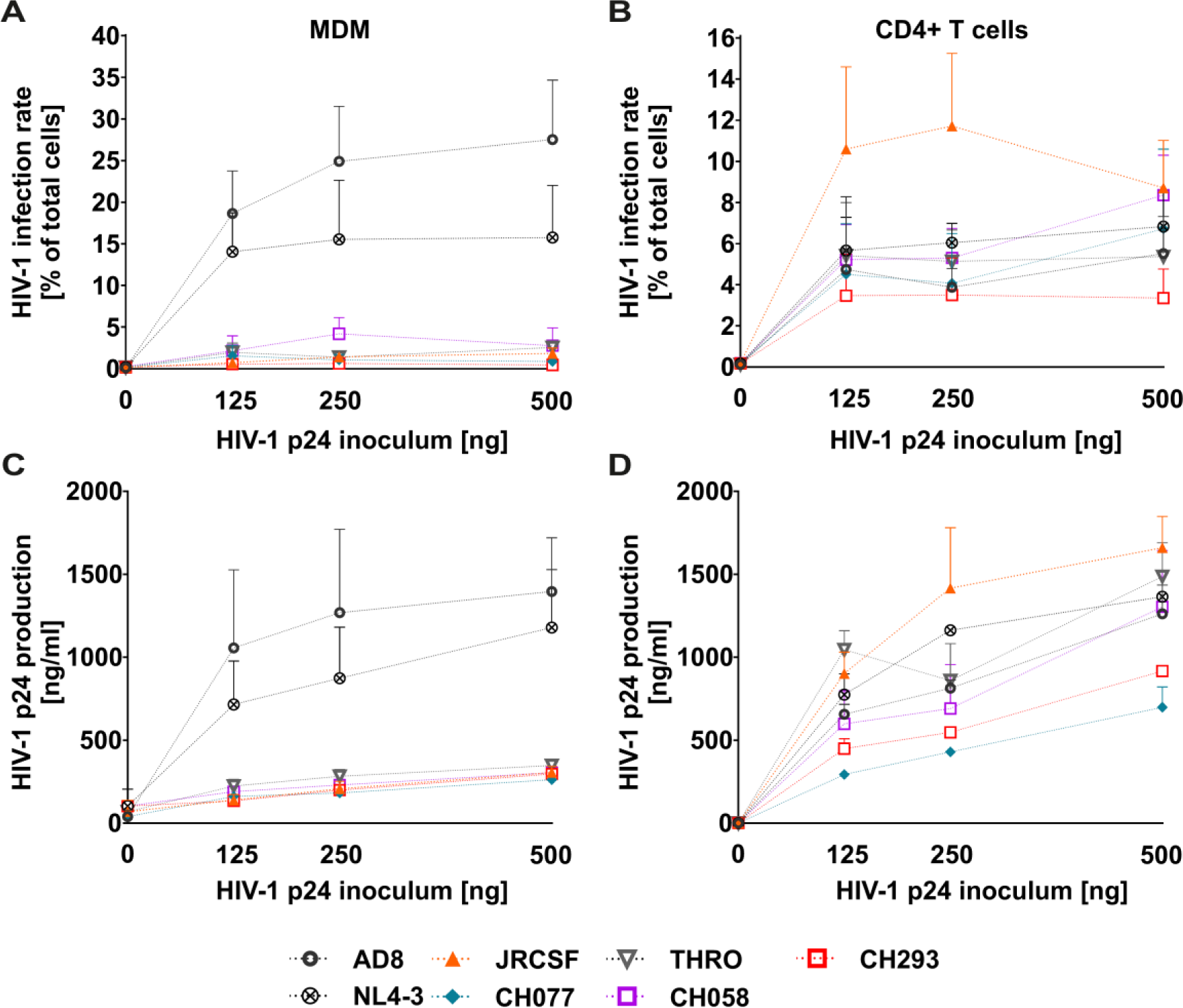
Primary HIV-1 strains replicate in CD4+ T cells but not in MDM. Monocyte derived macrophages (MDM) (A) or activated CD4+ T cells (B) were infected with 125, 250 or 500 ng of HIV-1 p24 protein (inoculum) for 16 h (A) or for 4 h (B). MDM were immunostained to detect HIV-1 p24 and DAPI and analyzed by fluorescence microcopy at 7 dpi. The CD4+ T cells were immunostained to detect HIV-1 p24 and analyzed by flow cytometry at 4 dpi. HIV-1 replication is shown as infection rate and expressed as percentage of p24+/DAPI+ macrophages (A) or CD4+ T cells (B), respectively. n=4 donors, +/− SD. (C) and (D) p24 HIV-1 protein amount in cell culture was evaluated by p24 ELISA in MDMs and CD4+ T cells culture supernatant at 7 dpi or at 4 dpi, respectively. n=4 donors, +/− SD.

In contrast, replication of all strains was readily detectable in CD4+ T cells analyzed at 4 days post infection (Fig. 1B). We chose this earlier time point as compared to macrophages, because there is spread and replication in the culture occurring prior to the onset of massive virus-induced cytotoxicity and activation-induced cell death (32, 33). In order to corroborate the measurement done on infected cells by IF, we assessed p24 production in cell culture supernatants over 7 days (macrophages, Fig. 1C) or 4 days (CD4+ T cells, Fig. 1D) of virus replication, which is an indicator for the overall efficiency of virus production in the cultures. Consistent with our previous observations, only lab-adapted AD8 and R5-tropic NL4-3 were efficiently released from infected macrophages, whereas all other strains showed no, to low signs of p24 production (Fig. 1C). In contrast, to CD4+ T cells sustained p24-release across all tested strains (Fig. 1D). In conclusion, while it was expected that the primary strains replicate in CD4+ T cells, albeit with differential efficacy, primary strains capable of utilizing CCR5 are strongly compromised in terms of macrophage infection and p24 production in this model.

### Coculture of HIV-1 infected macrophages with CD4+ T cells facilitates viral replication

Macrophages and CD4+ T cells interact via various mechanisms including direct cell-to-cell contacts as for instance the immunological synapse or indirectly by secretion of cytokines and chemokines. Overall, such non-virus specific interactions seem to promote virus replication and spread (34, 35). Furthermore, there is evidence for cytokine and chemokine secretion that might be both, beneficial as well as inhibitory in the context of virus replication (36–39). However, most studies in this area have focused on infected CD4+ T cells cocultured with uninfected macrophages (40–42). In contrast, especially for primary patient-derived HIV-1, less is known about overall effects of coculturing HIV-1 infected macrophages with uninfected CD4+ T cells. To analyze this, we infected macrophages with the different HIV-1 laboratory-adapted and primary strains using increasing amounts of virus normalized to the amount of p24. After 16 h, macrophages were washed to remove virus inoculum. Three days later, activated CD4+ T cells were added and left in coculture for additional 4 days. Then, after removal of the CD4+ T cells in the supernatants and excessive washing, we assessed the rates of macrophage infection and replication by p24 staining and counting of infected cells via automated medium-throughput fluorescence microscopy. CD4+ T cell infection was quantified via p24 staining and flow cytometry. Overall levels of virus replication and HIV-1 production in the cocultures were measured in supernatants by p24 ELISA.

Strikingly, coculturing the HIV-1 challenged macrophages with CD4+ T cells strongly enhanced levels of virally infected macrophages in the cultures (Fig. 2A). R5-tropic lab-adapted strains AD8 and NL4-3 more than doubled the productive infection rate of macrophages (∼25% without CD4+ T cells to ∼50% with CD4+ T cells for AD8 and for NL4-3 from ∼15% to ∼45% with CD4+ T cells; compare Fig. 1A to Fig. 2A). Throughout, for the primary strains the effect was even more dramatic. For instance, the HIV-1 T/F strain CH077 infected ∼20% of macrophages at 500 ng of p24 inoculum when CD4+ T cells were added (Fig. 2A) as compared to nearly no infection without addition of CD4+ T cells (Fig. 1A). We made similar observations for the T/F strain THRO and CH058 and the chronic HIV-1 strain CH293 (Fig. 2A as compared to Fig. 1A). Similarly, JRCSF achieved high infection rates of about 30% at 500 ng p24 in the inoculum (Fig. 2A), whereas the isolate was unable to replicate in macrophages without CD4+ T cell coculture (Fig. 1A). Of note, the number of cells scoring p24-positive in the cocultured CD4+ T cells largely mirrored the kinetics and differences observed in macrophages (Fig. 2B), albeit with lower overall infection rates that did not exceed replication levels when CD4+ T cells were infected and cultured alone (compare to Fig. 1B). Finally, p24-levels in the cell culture supernatants correlated with the number of p24+ cells in the coculture (Fig. 2C as compared to Fig. 2A and B), indicating that what we measure is indeed productive infection resulting in robust p24-release. As expected, levels of p24 released into the supernatants were also increased as compared to CD4+ T cells and macrophages cultured alone.

**Figure 2.**
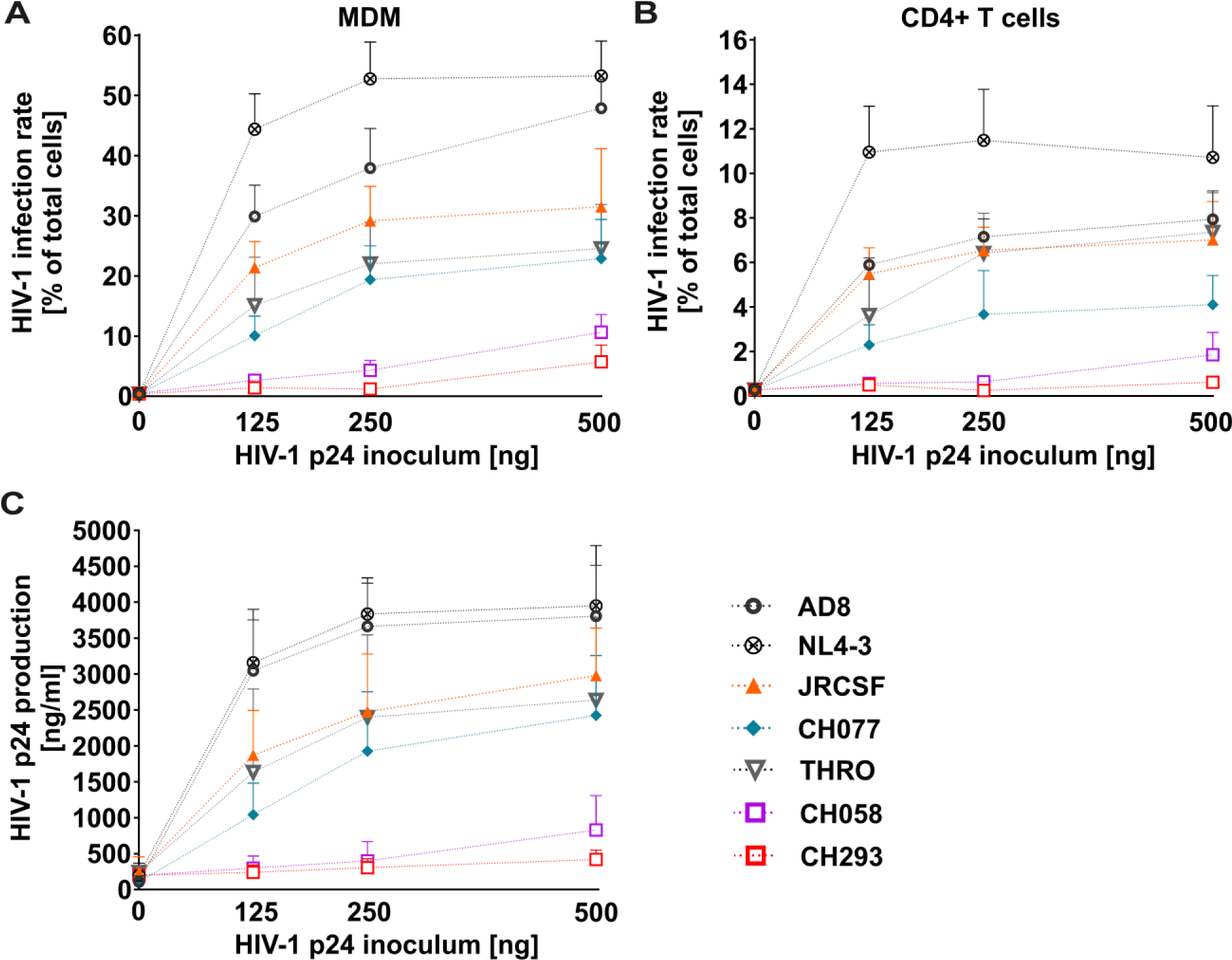
CD4+ T cells facilitate replication of chronic and T/F isolates in MDM. Monocyte derived macrophages (MDM) were infected with 125, 250 or 500 ng of HIV-1 p24 (inoculum) for 16 h, washed and subsequently activated CD4+ T cells were added at 3 dpi at a ratio of 1:10 (1 MDM: 10 CD4+ T cells). MDM were immunostained to detect HIV-1 p24 and counterstained with DAPI, followed by analysis via fluorescence microcopy at 7 dpi (A). The CD4+ T cells from the coculture were immunostained to detect HIV-1 p24 and analyzed by flow cytometry at 4 dpi by (B). HIV-1 replication is shown as infection rate and expressed as percentage of p24+/DAPI+ macrophages (A) or CD4+ T cells (B), respectively. p24 viral protein concentration was evaluated by p24 ELISA in coculture supernatant at 7dpi (C). n=4 donors, +/− SD.

Altogether, primary patient-derived HIV-1 strains, which are severely impaired in their ability to replicate in macrophages, efficiently spread in these cells upon coculture with non-infected CD4+ T cells.

### CD4+ T cells induce the formation of VCCs in HIV-1 infected MDM, correlating with virus replication

HIV-1 infection of macrophages leads to the sequestration of newly formed viruses in VCCs, variously characterized as a source of virus for trans-infection, a site of virus assembly, or a site of virus storage following assembly on the plasma membrane (12, 43–46). However, up to now, VCC formation was exclusively studied in lab-adapted strains NL4-3 and AD8 (12, 14, 15, 44–49). Hence, we first evaluated the ability of primary HIV-1 strains to form VCCs in macrophages and found, in agreement with the inability of these strains to seed productive infection in macrophages (Fig. 1A), few to no VCCs when we stained for intracellular p24 (Fig. 3). We then hypothesized that coculture of infected macrophages with non-infected CD4+ T cells might facilitate formation of VCCs. To address this, HIV-1 infected monocyte-derived macrophages (MDMs), cocultured with non-infected CD4+ T cells were stained against p24 in order to visualize the amount of VCCs and their subcellular distribution. In comparison to macrophages infected and cultured alone, we indeed observed the appearance of VCCs in macrophages infected with primary HIV-1 strains upon coculture with CD4+ T cells, even though they seemed fewer and less prominent compared to VCCs formed by lab-adapted strains (Fig. 3). When we quantified the total VCC area normalized to cell numbers (by counting the DAPI+ nuclei), the previous qualitative assessment became strikingly clear (Fig. 4A). Indeed, neither JRCSF, nor one of the other T/F and chronic strains were able to establish VCCs in MDM, whereas coculture initiated VCC formation. In contrast, the amount of VCCs was nearly identical when MDM were initially infected with AD8 or NL4-3 irrespective of adding non-infected CD4+ T cells to the cultures or not (Fig. 4A). Overall, coculturing HIV-1 infected MDM with non-infected CD4+ T cells robustly induced the formation of VCCs (Fig. 4B).

**Figure 3.**
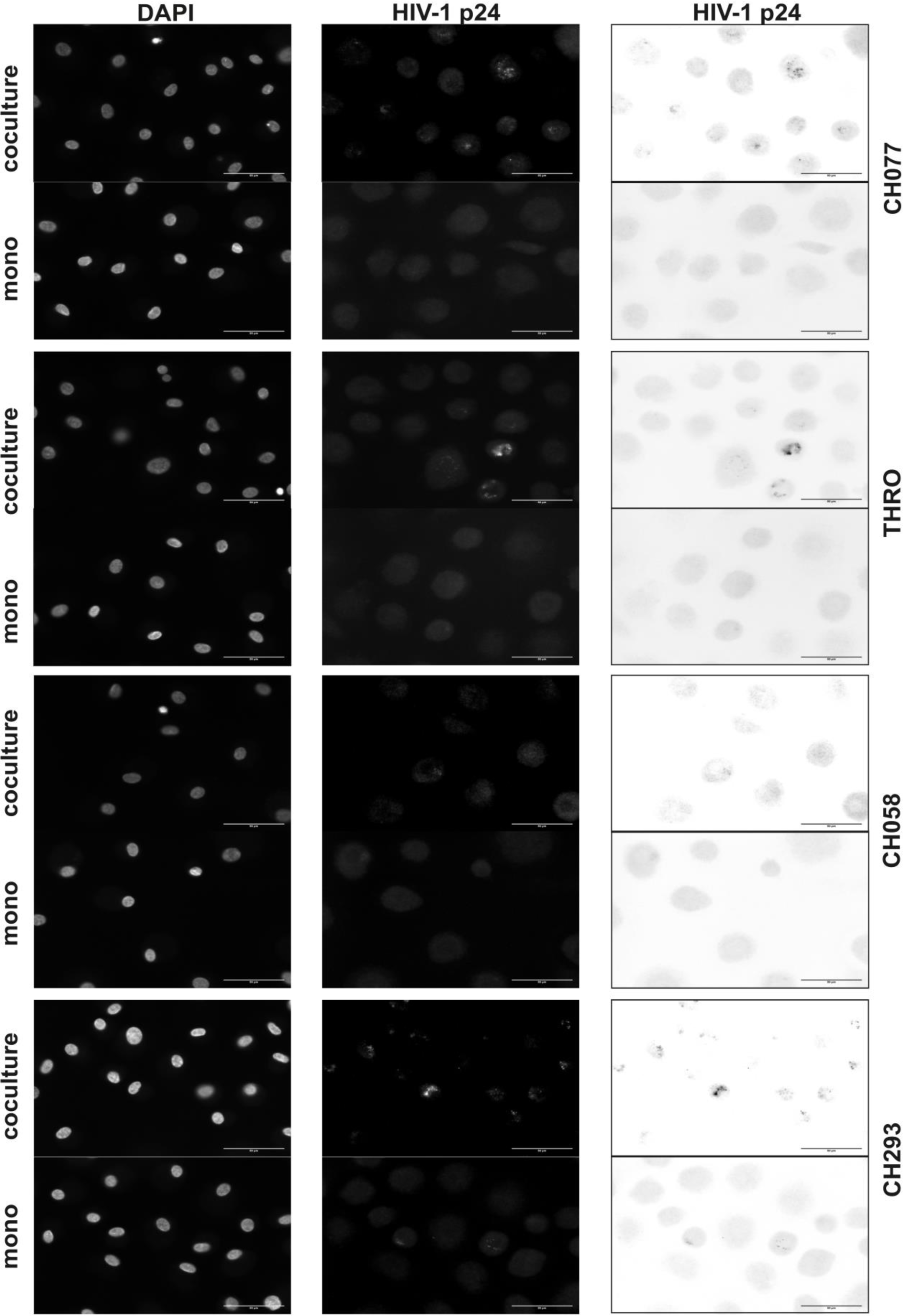
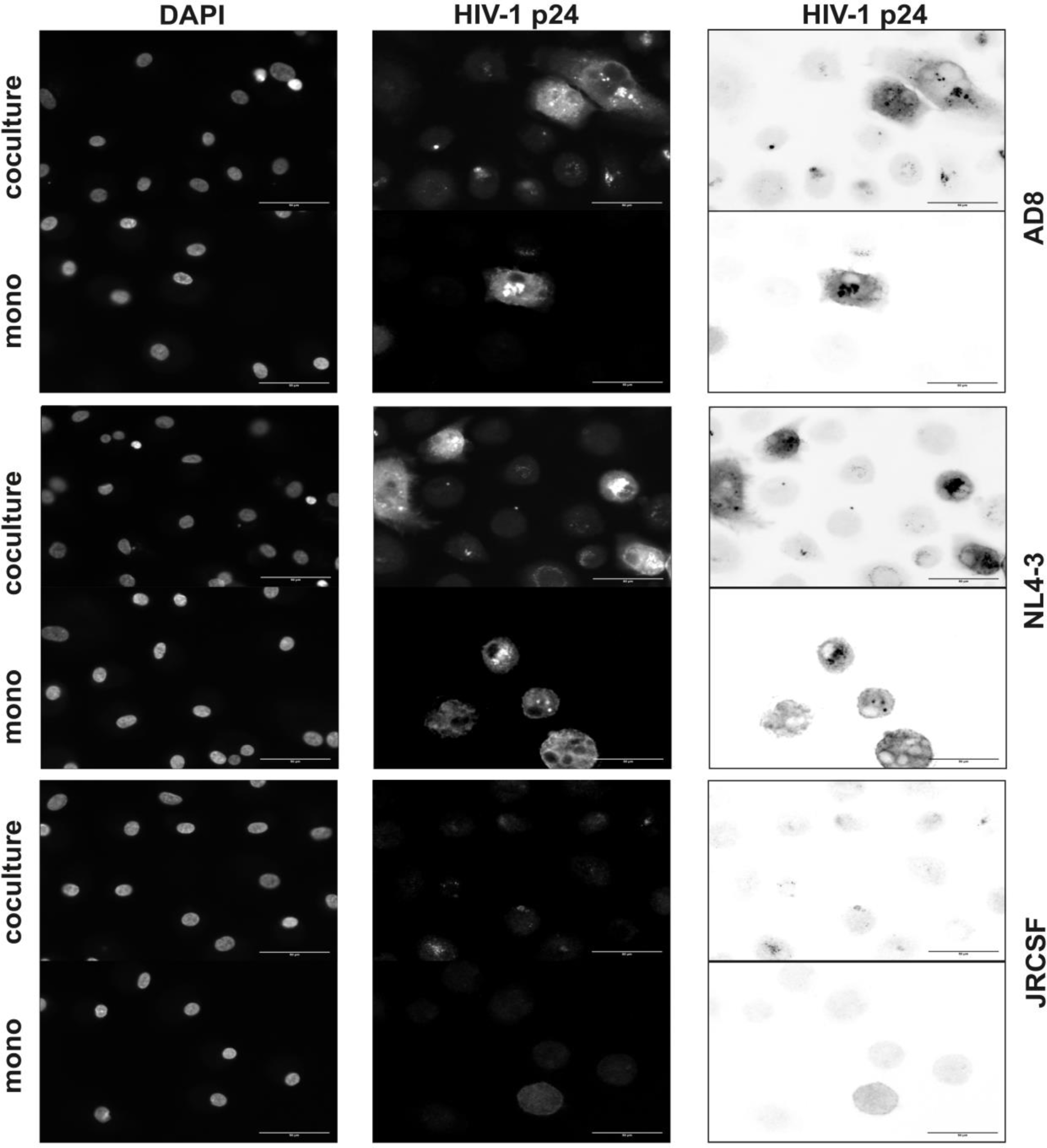
Formation of VCCs in HIV-1 infected MDM with or without CD4+ T cell coculture. Monocyte derived macrophages (MDM) were infected with 250 ng of HIV-1 p24 protein for 16 h, washed and subsequently activated CD4+ T cells were added at 3 dpi in a ratio of 1:10 (1 MDM: 10 CD4+ T cells). At 7 dpi MDM were immunostained to detect HIV-1 p24 and DAPI and analyzed by fluorescence microcopy. 40X magnification objective was used to analyze VCCs formation in the cultures. DAPI was used to identify nuclei and individual cells; p24 staining to detect infected cells and VCCs is shown as regular and inverted channel. The scale bar in the images represents 50 µm. The images are representative from one donor out of three donors for CH058c; 4 donors for AD8, JRCSF, NL4-3, CH077, THRO and CH293.

**Figure 4.**
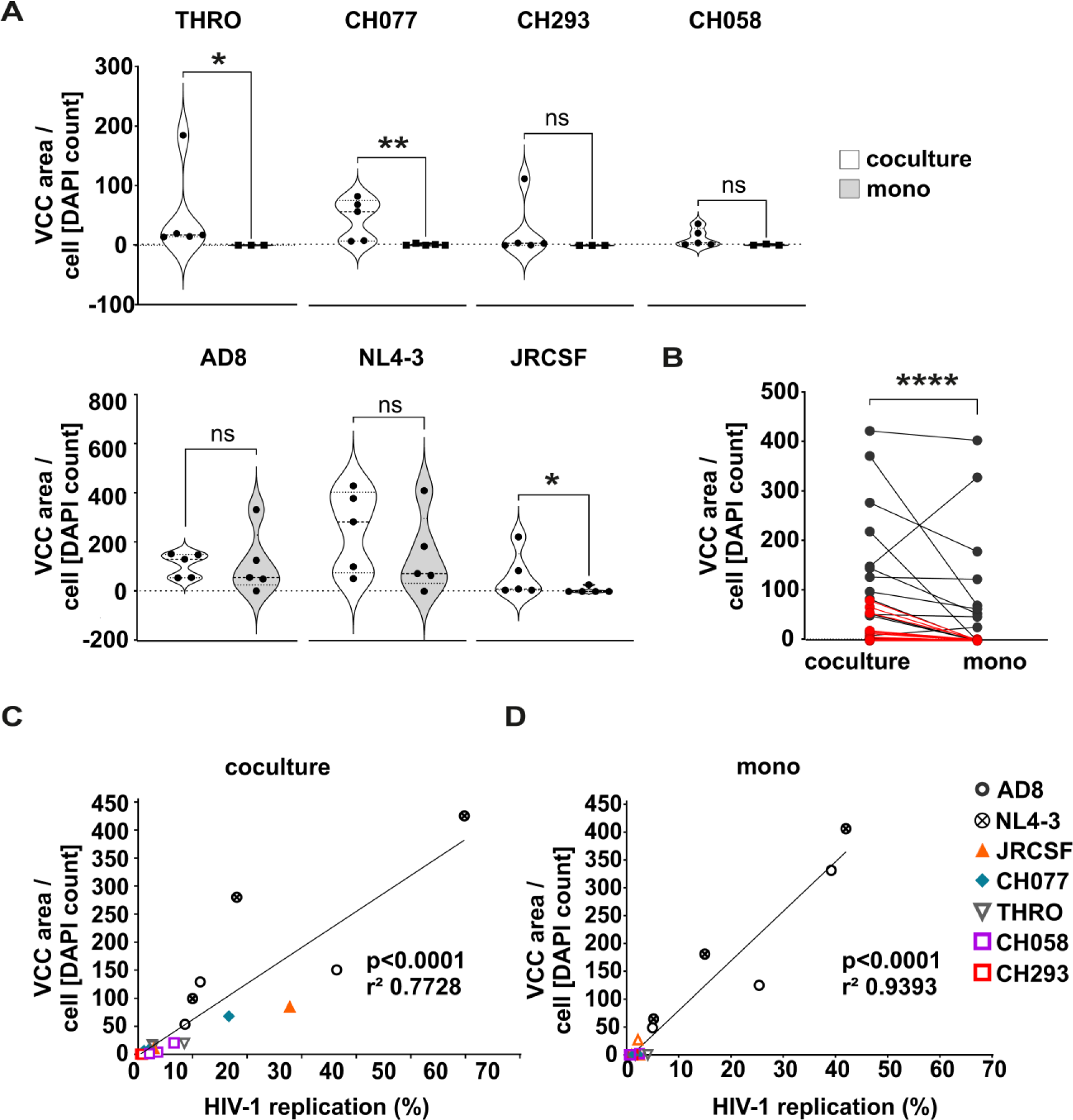
Non-infected CD4+ T cells induce formation of VCCs in HIV-1 infected MDM which correlates with virus replication. Monocyte derived macrophages (MDM) were infected with 250 ng of HIV-1 p24 protein for 16 h, washed and subsequently activated CD4+ T cells were added at 3 dpi in a ratio of 1:10 (1 MDM: 10 CD4+ T cells). At 7 dpi MDM were immunostained to detect HIV-1 p24 followed by counterstaining with DAPI and analysis via fluorescence microcopy at 40X magnification. (A) Calculation of VCCs area sum over DAPI counting in MDM from co-culture system (white violins) and from MDM cultured alone (grey violins) considering each HIV-1 strain. 5 donors for coculture system for all virus strains, for monoculture system 3 donors of CH058, THRO and Ch293 and 5 donors of AD8, JRCSF, NL4-3 and CH077. Significance was tested with unpaired T-test and Mann-Whitney test. *, p<0.05; **, p<0.01; ns, not significant. (B) The ratio VCCs area sum over DAPI counting is expressed comparing each system (coculture and monoculture) independently of the HIV-1 strain (black spots represent laboratory adapted strains and red spots represent the primary strains, n = 21 i.e. three biological replicates of each of the seven HIV-1 strains). Significance was tested with paired T-test and Wilcoxon matched-pairs signed rank test. ****, p<0.0001. C and D) Correlation between the VCCs area sum over DAPI counting and infection rate (expressed as HIV-1 replication rate in percentage) in MDM from co-culture (C) and MDM cultured alone (D). n=3 donors. The trendline shows a linear regression analysis with squared Pearson correlation coefficient (r^2^) and calculated statistical significance.

As previously mentioned, VCCs may function as storage compartments of HIV-1 in macrophages, or play more direct roles in HIV-1 trans-infection or spread. Given that coculture induced replication of primary strains in MDM as well as VCC formation, we hypothesized that these two phenotypes might be interconnected. Indeed, the VCC area correlated with HIV-1 replication rate in all the T/F strains (THRO, CH058, CH077), the HIV-1 chronic strain (CH293) as well as the lab adapted ones (AD8, JR-CSF, NL4-3) either in MDM from cocultures or MDM cultured alone (see Fig. 4 C and D, respectively). Altogether, replication of primary HIV-1 strains in MDM is strongly increased by coculture with non-infected CD4+ T cells, indicating that these strains depend on cellular cross-talk for efficient transmission and spread. Furthermore, non-infected CD4+ T cell coculture induces VCC formation in MDM and the abundance of VCCs correlates with virus replication.

## DISCUSSION

A major challenge in HIV-1 therapy and cure approaches is the management of long-term viral persistence in cells that act as a reservoir of the virus, including CD4+ memory T cells and tissue-resident macrophages. Hence, there is a strong interest in understanding modes of viral transmission and reactivation of virus replication in those cells (50–52). On top, it is essential to include primary HIV-1 isolates, such as T/F or chronic HIV-1 strains alongside different lab-adapted strains in such studies.

Ample previous studies have assessed how these isolates replicate in CD4+ T cells and macrophages and found in accordance to our data that T/F viruses are inefficient in infecting macrophages, even though they are capable of using CCR5 as co-receptor. However, their replication is readily detectable in CD4+ T cells (21, 35, 41). This block in macrophage infection can be overcome by cell-to-cell spread, where HIV-1 is directly transferred from macrophages to CD4+ T cells (16, 17, 35, 53–56). In addition, through processes like trans-cytosis, antigen-presenting cells including macrophages and DCs capture infectious virus to pass it on for CD4+ T cell infection (57). Also, the cellular environment generated after macrophage infection with lab-adapted or T/F HIV strains can enhance infection of resting CD4+ T cells (19) and can even skew the differentiation of activated CD4+ T cells into more permissive profiles (20). Vice versa, it is well established that HIV-1 infected CD4+ T cells transmit HIV-1 to macrophages through the virological synapse, direct cell-to-cell contact and even via phagocytosis and fusion with infected CD4+ T cells (56, 58–60).

In contrast, we are not aware of studies that investigate how HIV-1 infection in macrophages is influenced by coculture with autologous non-infected CD4+ T cells (56). Our findings highlight, that lab-adapted strains replicate efficiently in macrophages independent of CD4+ T cell coculture, whereas replication of primary HIV-1 strains in macrophages is recovered and strongly fueled by the addition of non-infected CD4+ T cells. Currently it is unclear if this enhancement of HIV-1 replication in macrophages is mediated by direct cellular interactions, or via secreted factors, or a combination of both and it will be important to assess this in future experiments.

Regardless, it became clear in our experiments that viral replication in macrophages is intimately linked to formation of macrophage internal virus-containing compartments (VCCs). VCCs are vacuolar subcellular structures that can be connected to the plasma membrane where it has been described that HIV-1 accumulates as infectious virions (12, 14, 15, 48, 49, 61). Macrophage internal HIV-1 in VCCs is shielded from recognition by broadly neutralizing antibodies (15, 62) and VCCs are characterized as storage and virus transmission compartments (8, 16, 47, 48). Here, we extended previous results showing the presence of VCCs in macrophages infected with not only lab-adapted HIV-1 strains, but also with T/F and chronic strains, which had not been studied before.

Our data indicate that indeed VCC formation correlates with the presence of CD4+ T cells in the coculture and notoriously increased virus replication in macrophages. Intriguingly, this implies that HIV-1 replication in macrophages may remain in an inactive silent state and will be fueled by the presence of non-infected activated CD4+ T cells. Given the fact that VCCs in macrophages seem to represent an immune-privileged niche refractory to neutralizing antibodies (15, 62), highlights the necessity to purge this reservoir through therapeutic intervention (50, 63). Of note, evidence suggests that virus release from macrophages is impaired, when VCC-anchored actin filaments are disrupted via experimental drugs (64).

In summary, our results suggest that patient-derived HIV-1 strains form VCCs in macrophages upon coculture with autologous non-infected T cells and that this process correlates with virus replication. Considering the important role of macrophages as reservoir cells, it will be essential to develop strategies that target HIV-1 in VCCs or destroy VCCs in conjunction with therapeutic interventions that aim to reduce the burden of latent HIV-1 in CD4+ T cells. Only by comprehensively addressing HIV-1 persistence in the variety of potential reservoirs, the nowadays set ambitious goal of HIV-1 functional cure might become achievable.

## Consent for publication

All authors gave their consent to publish. All authors read and approved the final manuscript.

## Availability of data and material

All data generated and analyzed during this study are included in this published manuscript.

## Authors’ contributions

SVM performed most of the experiments supported by JL and LMM. SVM and MS planned the experiments and analyzed the data. GT and MS provided resources and reagent tools. SVM and MS wrote the manuscript draft. MS supervised the overall study. All authors contributed to editing and developed the manuscript to its final form.

## Acknowledgements

We thank Tamam Bakchoul and the team of the Institute for Clinical and Experimental Transfusion Medicine (IKET) of the University Hospital Tuebingen for their constant support in providing buffy coats for preparation of primary CD4+ T cells and macrophages.

## REFERENCES

1. Barre-Sinoussi F, Chermann JC, Rey F, Nugeyre MT, Chamaret S, Gruest J, Dauguet C, Axler-Blin C, Vezinet-Brun F, Rouzioux C, Rozenbaum W, Montagnier L. 1983. Isolation of a T-lymphotropic retrovirus from a patient at risk for acquired immune deficiency syndrome (AIDS). Science 220:868–71.

2. www.who.int/data/gho/data/themes/hiv-aids. Accessed March 19th 2024.

3. Maartens G, Celum C, Lewin SR. 2014. HIV infection: epidemiology, pathogenesis, treatment, and prevention. Lancet 384:258–71.

4. Gulick RM, Mellors JW, Havlir D, Eron JJ, Gonzalez C, McMahon D, Richman DD, Valentine FT, Jonas L, Meibohm A, Emini EA, Chodakewitz JA. 1997. Treatment with indinavir, zidovudine, and lamivudine in adults with human immunodeficiency virus infection and prior antiretroviral therapy. N Engl J Med 337:734–9.

5. Rodger AJ, Lodwick R, Schechter M, Deeks S, Amin J, Gilson R, Paredes R, Bakowska E, Engsig FN, Phillips A. 2013. Mortality in well controlled HIV in the continuous antiretroviral therapy arms of the SMART and ESPRIT trials compared with the general population. Aids 27:973–9.

6. Deeks SG, Archin N, Cannon P, Collins S, Jones RB, de Jong M, Lambotte O, Lamplough R, Ndung’u T, Sugarman J, Tiemessen CT, Vandekerckhove L, Lewin SR, International ASGSSwg. 2021. Research priorities for an HIV cure: International AIDS Society Global Scientific Strategy 2021. Nat Med 27:2085–2098.

7. Coleman CM, Wu L. 2009. HIV interactions with monocytes and dendritic cells: viral latency and reservoirs. Retrovirology 6:51.

8. Koppensteiner H, Brack-Werner R, Schindler M. 2012. Macrophages and their relevance in Human Immunodeficiency Virus Type I infection. Retrovirology 9:82.

9. Janeway CA, Travers P, Walport M, Shlomchik MJ. 2005. Immunobiology: the immune system in health and disease.

10. Ganor Y, Real F, Sennepin A, Dutertre CA, Prevedel L, Xu L, Tudor D, Charmeteau B, Couedel-Courteille A, Marion S, Zenak AR, Jourdain JP, Zhou Z, Schmitt A, Capron C, Eugenin EA, Cheynier R, Revol M, Cristofari S, Hosmalin A, Bomsel M. 2019. HIV-1 reservoirs in urethral macrophages of patients under suppressive antiretroviral therapy. Nat Microbiol 4:633–644.

11. Woottum M, Yan S, Sayettat S, Grinberg S, Cathelin D, Bekaddour N, Herbeuval JP, Benichou S. 2024. Macrophages: Key Cellular Players in HIV Infection and Pathogenesis. Viruses 16.

12. Deneka M, Pelchen-Matthews A, Byland R, Ruiz-Mateos E, Marsh M. 2007. In macrophages, HIV-1 assembles into an intracellular plasma membrane domain containing the tetraspanins CD81, CD9, and CD53. J Cell Biol 177:329–41.

13. Wiley RD, Gummuluru S. 2006. Immature dendritic cell-derived exosomes can mediate HIV-1 trans infection. Proc Natl Acad Sci U S A 103:738–43.

14. Welsch S, Keppler OT, Habermann A, Allespach I, Krijnse-Locker J, Krausslich HG. 2007. HIV-1 buds predominantly at the plasma membrane of primary human macrophages. PLoS Pathog 3:e36.

15. Koppensteiner H, Banning C, Schneider C, Hohenberg H, Schindler M. 2012. Macrophage internal HIV-1 is protected from neutralizing antibodies. J Virol 86:2826–36.

16. Gousset K, Ablan SD, Coren LV, Ono A, Soheilian F, Nagashima K, Ott DE, Freed EO. 2008. Real-time visualization of HIV-1 GAG trafficking in infected macrophages. PLoS Pathog 4:e1000015.

17. Parrish NF, Gao F, Li H, Giorgi EE, Barbian HJ, Parrish EH, Zajic L, Iyer SS, Decker JM, Kumar A, Hora B, Berg A, Cai F, Hopper J, Denny TN, Ding H, Ochsenbauer C, Kappes JC, Galimidi RP, West AP, Jr., Bjorkman PJ, Wilen CB, Doms RW, O’Brien M, Bhardwaj N, Borrow P, Haynes BF, Muldoon M, Theiler JP, Korber B, Shaw GM, Hahn BH. 2013. Phenotypic properties of transmitted founder HIV-1. Proc Natl Acad Sci U S A 110:6626–33.

18. Dufloo J, Bruel T, Schwartz OJR. 2018. HIV-1 cell-to-cell transmission and broadly neutralizing antibodies. 15:51.

19. Trifone C, Salido J, Ruiz MJ, Leng L, Quiroga MF, Salomon H, Bucala R, Ghiglione Y, Turk G. 2018. Interaction Between Macrophage Migration Inhibitory Factor and CD74 in Human Immunodeficiency Virus Type I Infected Primary Monocyte-Derived Macrophages Triggers the Production of Proinflammatory Mediators and Enhances Infection of Unactivated CD4(+) T Cells. Front Immunol 9:1494.

20. Trifone C, Baquero L, Czernikier A, Benencio P, Leng L, Laufer N, Quiroga MF, Bucala R, Ghiglione Y, Turk G. 2022. Macrophage Migration Inhibitory Factor (MIF) Promotes Increased Proportions of the Highly Permissive Th17-like Cell Profile during HIV Infection. Viruses 14.

21. Ochsenbauer C, Edmonds TG, Ding H, Keele BF, Decker J, Salazar MG, Salazar-Gonzalez JF, Shattock R, Haynes BF, Shaw GM, Hahn BH, Kappes JC. 2012. Generation of transmitted/founder HIV-1 infectious molecular clones and characterization of their replication capacity in CD4 T lymphocytes and monocyte-derived macrophages. J Virol 86:2715–28.

22. Gramberg T, Sunseri N, Landau NR. 2010. Evidence for an activation domain at the amino terminus of simian immunodeficiency virus Vpx. J Virol 84:1387–96.

23. Businger R, Kivimaki S, Simeonov S, Vavouras Syrigos G, Pohlmann J, Bolz M, Muller P, Codrea MC, Templin C, Messerle M, Hamprecht K, Schaffer TE, Nahnsen S, Schindler M. 2021. Comprehensive Analysis of Human Cytomegalovirus- and HIV-Mediated Plasma Membrane Remodeling in Macrophages. mBio 12:e0177021.

24. Schindler M, Rajan D, Banning C, Wimmer P, Koppensteiner H, Iwanski A, Specht A, Sauter D, Dobner T, Kirchhoff F. 2010. Vpu serine 52 dependent counteraction of tetherin is required for HIV-1 replication in macrophages, but not in ex vivo human lymphoid tissue. Retrovirology 7:1.

25. Picchio GR, Gulizia RJ, Wehrly K, Chesebro B, Mosier DE. 1998. The cell tropism of human immunodeficiency virus type 1 determines the kinetics of plasma viremia in SCID mice reconstituted with human peripheral blood leukocytes. J Virol 72:2002–9.

26. Schubert U, Bour S, Willey RL, Strebel K. 1999. Regulation of virus release by the macrophage-tropic human immunodeficiency virus type 1 AD8 isolate is redundant and can be controlled by either Vpu or Env. J Virol 73:887–96.

27. Chavda SC, Griffin P, Han-Liu Z, Keys B, Vekony MA, Cann AJ. 1994. Molecular determinants of the V3 loop of human immunodeficiency virus type 1 glycoprotein gp120 responsible for controlling cell tropism. J Gen Virol 75 (Pt 11):3249–53.

28. Koyanagi Y, Miles S, Mitsuyasu RT, Merrill JE, Vinters HV, Chen IS. 1987. Dual infection of the central nervous system by AIDS viruses with distinct cellular tropisms. Science 236:819–22.

29. Hrecka K, Hao C, Gierszewska M, Swanson SK, Kesik-Brodacka M, Srivastava S, Florens L, Washburn MP, Skowronski J. 2011. Vpx relieves inhibition of HIV-1 infection of macrophages mediated by the SAMHD1 protein. Nature 474:658–61.

30. Laguette N, Sobhian B, Casartelli N, Ringeard M, Chable-Bessia C, Segeral E, Yatim A, Emiliani S, Schwartz O, Benkirane M. 2011. SAMHD1 is the dendritic- and myeloid-cell-specific HIV-1 restriction factor counteracted by Vpx. Nature 474:654–7.

31. Businger R, Deutschmann J, Gruska I, Milbradt J, Wiebusch L, Gramberg T, Schindler M. 2019. Human cytomegalovirus overcomes SAMHD1 restriction in macrophages via pUL97. Nat Microbiol 4:2260–2272.

32. Gandhi RT, Chen BK, Straus SE, Dale JK, Lenardo MJ, Baltimore D. 1998. HIV-1 directly kills CD4+ T cells by a Fas-independent mechanism. J Exp Med 187:1113–22.

33. Groux H, Torpier G, Monte D, Mouton Y, Capron A, Ameisen JC. 1992. Activation-induced death by apoptosis in CD4+ T cells from human immunodeficiency virus-infected asymptomatic individuals. J Exp Med 175:331–40.

34. Pedro KD, Henderson AJ, Agosto LM. 2019. Mechanisms of HIV-1 cell-to-cell transmission and the establishment of the latent reservoir. Virus Res 265:115–121.

35. Calado M, Pires D, Conceicao C, Santos-Costa Q, Anes E, Azevedo-Pereira JM. 2023. Human immunodeficiency virus transmission-Mechanisms underlying the cell-to-cell spread of human immunodeficiency virus. Rev Med Virol 33:e2480.

36. Vandergeeten C, Fromentin R, Chomont N. 2012. The role of cytokines in the establishment, persistence and eradication of the HIV reservoir. Cytokine Growth Factor Rev 23:143–9.

37. Catalfamo M, Le Saout C, Lane HC. 2012. The role of cytokines in the pathogenesis and treatment of HIV infection. Cytokine Growth Factor Rev 23:207–14.

38. Keating SM, Jacobs ES, Norris PJ. 2012. Soluble mediators of inflammation in HIV and their implications for therapeutics and vaccine development. Cytokine Growth Factor Rev 23:193–206.

39. Kedzierska K, Crowe SM. 2001. Cytokines and HIV-1: interactions and clinical implications. Antivir Chem Chemother 12:133–50.

40. Cong L, Sugden SM, Leclair P, Lim CJ, Pham TNQ, Cohen EA. 2021. HIV-1 Vpu Promotes Phagocytosis of Infected CD4(+) T Cells by Macrophages through Downregulation of CD47. mBio 12:e0192021.

41. Han M, Cantaloube-Ferrieu V, Xie M, Armani-Tourret M, Woottum M, Pages JC, Colin P, Lagane B, Benichou S. 2022. HIV-1 cell-to-cell spread overcomes the virus entry block of non-macrophage-tropic strains in macrophages. PLoS Pathog 18:e1010335.

42. Xie M, Leroy H, Mascarau R, Woottum M, Dupont M, Ciccone C, Schmitt A, Raynaud-Messina B, Verollet C, Bouchet J, Bracq L, Benichou S. 2019. Cell-to-Cell Spreading of HIV-1 in Myeloid Target Cells Escapes SAMHD1 Restriction. mBio 10.

43. Deneka M, Pelchen-Matthews A, Byland R, Ruiz-Mateos E, Marsh MJTJocb. 2007. In macrophages, HIV-1 assembles into an intracellular plasma membrane domain containing the tetraspanins CD81, CD9, and CD53. 177:329–341.

44. Raposo G, Moore M, Innes D, Leijendekker R, Leigh-Brown A, Benaroch P, Geuze H. 2002. Human macrophages accumulate HIV-1 particles in MHC II compartments. Traffic 3:718–29.

45. Pelchen-Matthews A, Kramer B, Marsh M. 2003. Infectious HIV-1 assembles in late endosomes in primary macrophages. J Cell Biol 162:443–55.

46. Hammonds JE, Beeman N, Ding L, Takushi S, Francis AC, Wang JJ, Melikyan GB, Spearman P. 2017. Siglec-1 initiates formation of the virus-containing compartment and enhances macrophage-to-T cell transmission of HIV-1. PLoS Pathog 13:e1006181.

47. Gaudin R, de Alencar BC, Jouve M, Berre S, Le Bouder E, Schindler M, Varthaman A, Gobert FX, Benaroch P. 2012. Critical role for the kinesin KIF3A in the HIV life cycle in primary human macrophages. J Cell Biol 199:467–79.

48. Gaudin R, Berre S, Cunha de Alencar B, Decalf J, Schindler M, Gobert FX, Jouve M, Benaroch P. 2013. Dynamics of HIV-containing compartments in macrophages reveal sequestration of virions and transient surface connections. PLoS One 8:e69450.

49. Jouve M, Sol-Foulon N, Watson S, Schwartz O, Benaroch P. 2007. HIV-1 buds and accumulates in “nonacidic” endosomes of macrophages. Cell Host Microbe 2:85–95.

50. Graziano F, Vicenzi E, Poli G. 2016. Immuno-Pharmacological Targeting of Virus-Containing Compartments in HIV-1-Infected Macrophages. Trends Microbiol 24:558–567.

51. Rodrigues V, Ruffin N, San-Roman M, Benaroch P. 2017. Myeloid Cell Interaction with HIV: A Complex Relationship. Front Immunol 8:1698.

52. Ikeogu N, Ajibola O, Zayats R, Murooka TT. 2023. Identifying physiological tissue niches that support the HIV reservoir in T cells. mBio 14:e0205323.

53. van Teijlingen NH, Eder J, Sarrami-Forooshani R, Zijlstra-Willems EM, Roovers JWR, van Leeuwen E, Ribeiro CMS, Geijtenbeek TBH. 2023. Immune activation of vaginal human Langerhans cells increases susceptibility to HIV-1 infection. Sci Rep 13:3283.

54. Lopez P, Koh WH, Hnatiuk R, Murooka TT. 2019. HIV Infection Stabilizes Macrophage-T Cell Interactions To Promote Cell-Cell HIV Spread. J Virol 93.

55. Groot F, Welsch S, Sattentau QJ. 2008. Efficient HIV-1 transmission from macrophages to T cells across transient virological synapses. Blood 111:4660–3.

56. Dupont M, Sattentau QJ. 2020. Macrophage Cell-Cell Interactions Promoting HIV-1 Infection. Viruses 12.

57. Shen R, Kappes JC, Smythies LE, Richter HE, Novak L, Smith PD. 2014. Vaginal myeloid dendritic cells transmit founder HIV-1. J Virol 88:7683–8.

58. Mascarau R, Woottum M, Fromont L, Gence R, Cantaloube-Ferrieu V, Vahlas Z, Leveque K, Bertrand F, Beunon T, Metais A, El Costa H, Jabrane-Ferrat N, Gallois Y, Guibert N, Davignon JL, Favre G, Maridonneau-Parini I, Poincloux R, Lagane B, Benichou S, Raynaud-Messina B, Verollet C. 2023. Productive HIV-1 infection of tissue macrophages by fusion with infected CD4+ T cells. J Cell Biol 222.

59. Han M, Woottum M, Mascarau R, Vahlas Z, Verollet C, Benichou S. 2022. Mechanisms of HIV-1 cell-to-cell transfer to myeloid cells. J Leukoc Biol 112:1261–1271.

60. Bracq L, Xie M, Lambele M, Vu LT, Matz J, Schmitt A, Delon J, Zhou P, Randriamampita C, Bouchet J, Benichou S. 2017. T Cell-Macrophage Fusion Triggers Multinucleated Giant Cell Formation for HIV-1 Spreading. J Virol 91.

61. Benaroch P, Billard E, Gaudin R, Schindler M, Jouve M. 2010. HIV-1 assembly in macrophages. Retrovirology 7:29.

62. Chu H, Wang JJ, Qi M, Yoon JJ, Wen X, Chen X, Ding L, Spearman P. 2012. The intracellular virus-containing compartments in primary human macrophages are largely inaccessible to antibodies and small molecules. PLoS One 7:e35297.

63. Berre S, Gaudin R, Cunha de Alencar B, Desdouits M, Chabaud M, Naffakh N, Rabaza-Gairi M, Gobert FX, Jouve M, Benaroch P. 2013. CD36-specific antibodies block release of HIV-1 from infected primary macrophages and its transmission to T cells. J Exp Med 210:2523–38.

64. Rodrigues V, Taheraly S, Maurin M, San-Roman M, Granier E, Hanouna A, Benaroch P. 2022. Release of HIV-1 particles from macrophages is promoted by an anchored cytoskeleton and driven by mechanical constraints. J Cell Sci 135.

